# Comparative genomics between Colombian clinical isolates of Monophasic Variant *Salmonella* Typhimurium and international clonal lineages

**DOI:** 10.1101/2023.06.23.546254

**Authors:** Cuenca-Arias Paloma, Montaño Lucy Angeline, Rodriguez Edna Catering, Ruiz-Moreno Héctor Alejandro, Montilla-Escudero Efraín, Villarreal José Miguel, Wiesner Magdalena

**Affiliations:** Grupo de Microbiologia, Instituto Nacional de Salud, Bogotá, Colombia; Chemistry Department, Faculty of Sciences, Universidad Nacional de Colombia, Ciudad Universitaria, Carrera 30 N° 45-03, 111321, Bogotá, Colombia; Grupo de Genómica de Microorganismos Emergentes. Dirección de Investigación en Salud Pública, Instituto Nacional de Salud, Bogotá, Colombia

**Keywords:** *Salmonella* Typhimurium, monophasic variant, Acute Diarrheal Disease, antimicrobial resistance, whole-genome sequencing, genomic surveillance, clonal lineages

## Abstract

In Colombia, *Salmonella* Typhimurium monophasic variant (MVST) is the fourth serovar recovered in laboratory surveillance of acute diarrheal disease (ADD). Given its rapid worldwide dissemination, increasing multidrug-resistance, and the emergence of different endemic clones, it is considered an emerging public health problem. This study compared 21 Colombian clinical isolates and 27 MVST genomes from Europe, Asia, the United States, and Australia to know the gene pool and to define similarities with international clones. Eighty percent of the Colombian MVST isolates formed a lineage divided into 2 clones, while 4 genomes were associated with the European ST34 and USA lineages. These two Colombian clones emerged in relatively recent events, in which possible spread was established during 2011 and 2012, exhibiting a diversity of plasmids and prophages, adapting to the Colombian population after differentiation. These results are a clear example of the high plasticity of MVST, evidencing the need for active genomic surveillance to monitor the circulation of new clonal lineages.

## Introduction

Salmonellosis is one of the most common foodborne diseases, with an estimated 95.1 million cases of acute gastroenteritis per year globally (1), it is caused for *Salmonella* spp., classified as more than 2600 serovars associated with both animals and humans and has a high impact on the economy and public health (2, 3).

Serovars are classified by the agglutination of antigens found on the bacterial cell wall (O antigen) and flagella (H antigen), flagellar antigens have 2 phases (phase 1 and phase 2 antigens) according to the specific combination of antigens a name is assigned to each serovar (3, 4). The main serovar associated with cases of human salmonellosis is *Salmonella* Typhimurium (Typhimurium), characterized by the successive appearance of dominant clones, including a monophasic variant lacking the second flagellar phase *fljAB* operon, which emerged in the late 1990s in Europe and the United States (USA) (5, 6) In the last 2 decades, *Salmonella* Typhimurium Monophasic Variant (MVST) has spread rapidly worldwide, displacing Typhimurium according to enteric disease surveillance reports (7). In MVST, the random loss of regions along the chromosome, the acquisition of resistance elements to antibiotics and heavy metals, as well as the differentiation between multidrug resistance (MDR), sequenciotype (ST), and phagetype profiles suggest their appearance by multiple independent events of source or geographic origin, giving rise to endemic clone (8, 9). Three primary clones stand out: European, Spanish, and USA clones, the first two, completely lack the *fljAB* operon, are MDR, and differ to ST34 or ST19, the absence of the virulence plasmid and acquisition of genomic island 4 (SGI-4); while the USA clone retains a *fljAB* operon gene and is antibiotic susceptible (10, 11).

In Colombia, the laboratory surveillance program for acute diarrheal disease (ADD), led by the Microbiology Group of the Instituto Nacional de Salud (INS), has carried out characterization to *Salmonella* spp. isolates recovered from clinical samples in the country for more than 20 years. In the year 2015, the circulation of MVST was reported for the first time in surveillance reports, and from that report until 2020, a total of 309 isolates of the monophasic variant were confirmed, positioning it serovar among the 4 most frequently (12, 13). Colombian isolates of MVST totally or partially lack the *fljAB* operon, exhibit resistance to tetracycline (TET), chloramphenicol (CHL), nalidixic acid (NAL), and ampicillin (AMP), and were preliminarily related to the European, Spanish, and USA clonal lineages. The circulation of the single-phase epidemic clone associated with the European ST34 clonal lineage was confirmed in 2019 (14, 15). This study aims to define the lineage of MVST isolates circulating in Colombia, as well as their genomic characteristics, for which comparative genomics was performed between 21 sequences of Colombian MVST clinical isolates and 27 MVST genomes from various sources, from Europe, Asia, USA, and Australia taking into account the clonal lineage of the isolates. As a result, we observed the circulation of two global MVST lineages in Colombia, and the inference of two endemic clonal groups in the country.

## Materials and Methods

### Colombian MVST clinical isolates

#### Colombian clinical isolates of MVST

For this study, 21 complete genomes of clinical isolates of MVST, recovered in Colombia from 2015 to 2018, were analyzed (Table supp1). Seven isolates were randomly selected from those previously analyzed (14), for whole genome sequencing (WGS), recovered from 2015 to 2018, have serotyping by the Kauffman-White-Le minor scheme (4), confirmed as single-phase Typhimurium serovar by PCR and susceptibility profiling (14). The antimicrobials evaluated were ampicillin, chloramphenicol, streptomycin, tetracycline, gentamicin, amikacin, nalidixic acid, trimethoprim, ciprofloxacin, ceftazidime, cefotaxime, by the Kirby Bauer method according to CLSI recommendations (16). To complement the study, 14 Colombian MVST sequences previously reported by Li et al (2019) that have all the phenotypic information previously mentioned were included (15).

Genomic DNA was obtained with the PureLink genomic DNA mini kit. Libraries were prepared with the Nextera XT DNA sample preparation kit (17). Sequencing was performed on a MiSeq system (Illumina). Reads were assembled with Trimmomatic v0.39 (https://github.com/timflutre/trimmomatic) and SPAdes v3.14.0 (https://github.com/ablab/spades) using default parameters.

### Selection of international MVST genomes

Twenty-seven international MVST sequences reported in GenBank from 1997 to 2018 were included, for 48 genomes analyzed. A search was performed in EMBL-EBI and PubMed databases, using keywords such as "phylogeny", "monophasic *Salmonella* Typhimurium ", "clonal lineage", "spain clone", "USA clone", "Europe clone", and geographical location, year of collection and source (human, food or animal) were considered as selection criteria (18–21). Twenty-seven genomes were selected from different sources and countries: United Kingdom (n = 1), USA (n = 15), Australia (n = 1), Spain (n = 3), and Japan (n = 7) (Table supplementary 1). MVST clones of Spanish (Accession number: DRR106818), European (Accession number: ERR985368), and USA (Accession number: ABAO01000000) origin were used as reference genomes together with the Typhimurium LT2 genome (Accession number: AE006468) in all analyses.

### Genomic Analysis

The 48 genomes selected for this study (21 Colombian MVSTs and 27 international MVSTs) were assembled and annotated with the BV-BRC: Bacterial and Viral Bioinformatics Resource Center (https://www.bv-brc.org) (22).

Several of the programs hosted on the Center for Genomic Epidemiology web platform were used (http://www.genomicepidemiology.org/), such as the MLST web tool (23) and the CSIPhylogeny-1.4 program for phylogeny from SNPs (24) using the default conditions (https://cge.cbs.dtu.dk/services/CSIPhylogeny/): minimum depth at SNP positions: 10, relative depth at SNP positions: 10, a minimum distance between SNPs: 10, minimum SNP quality: 30, minimum read mapping quality: 25, minimum Z-score: 1.96.

PlasmidFinder was used to determine plasmid incompatibility groups (25). For mass screening of antimicrobial genes, the ABRicate tool available on the Galaxy web platform Version 1.0.1 (https://usegalaxy.org/) was used by matching the ARG-ANNOT, CARD, NCBI bacterial Antimicrobial Resistance Reference Gene Database, Resfinder, the virulence gene detection the VFDB database was used. Proksee (https://proksee.ca) provides was used to visualize plasmid alignments (26).

The PHASTER (PHAge Search Tool Enhanced Release) program (http://phaster.ca/) was used to identify the prophage regions; the results of prophages that were intact or questionable were taken into account as presence.

For the pangenome, the PROKKA program was used, whose result in GFF3 format is used as input in the ROARY program (programs available in Galaxy Version 1.0.1 (https://usegalaxy.org/). Visualization was performed with the PHANDANGO program.

### Bayesian analysis - Time analysis in phylogeny

A maximum likelihood phylogeny was constructed using IQTREE v.1.6.12 (27) with the GTR+F+ASC+G4 substitution model chosen according to the Bayesian information criterion (BIC) (28). To understand the phylodynamic of MVST samples and estimate divergence dates, a Bayesian phylogenetic analysis was performed employing BEAST2 v.2.6.7 (29) defining sample dates as the date of leaves to calibrate the molecular clock. Preliminary studies were performed with the Nested Sampling package (15, 30, 31) for different molecular clock models and three prior probabilities. Strict, Relaxed Log-Normal and Random clocks were tested, along with Constant Coalescent, Exponential Coalescent, or Bayesian Coalescent (BSP) population. For all combinations of models and states a string size of 100000 steps was used with sampling every 5000 steps and burn-in of the first 100000 steps. A Relaxed Log Normal molecular clock model with a constant coalescent population was chosen according to the marginal likelihood value. The final analysis employed a chain size of 10000000000 steps with sampling every 5000 steps and burn-in of the first 1000000 steps. The phylogeny was visualized in FigTree v1.4.4 (http://tree.bio.ed.ac.uk/software/figtree/).

## Results

### Eighty percent of the Colombian isolates belong to two endemic clones

To know the phylogenetic relationship of the Colombian genomes, a tree was constructed from the variations in the genome using the maximum likelihood method. Of the 48 genomes analyzed, four sequences were identified and confirmed. The ancestral ST19 was the most common, found in 31 isolates. The European ST34 was found in 12 isolates, the ST2379 was found in one isolate reported in the USA, and the new ST7478 was found in two Colombian isolates. The ST7478 sequence differs from ST19 in the *sucA* allele. This study is the first to report the association of ST7478 with MVST (Figure 1).

**Figure 1.**
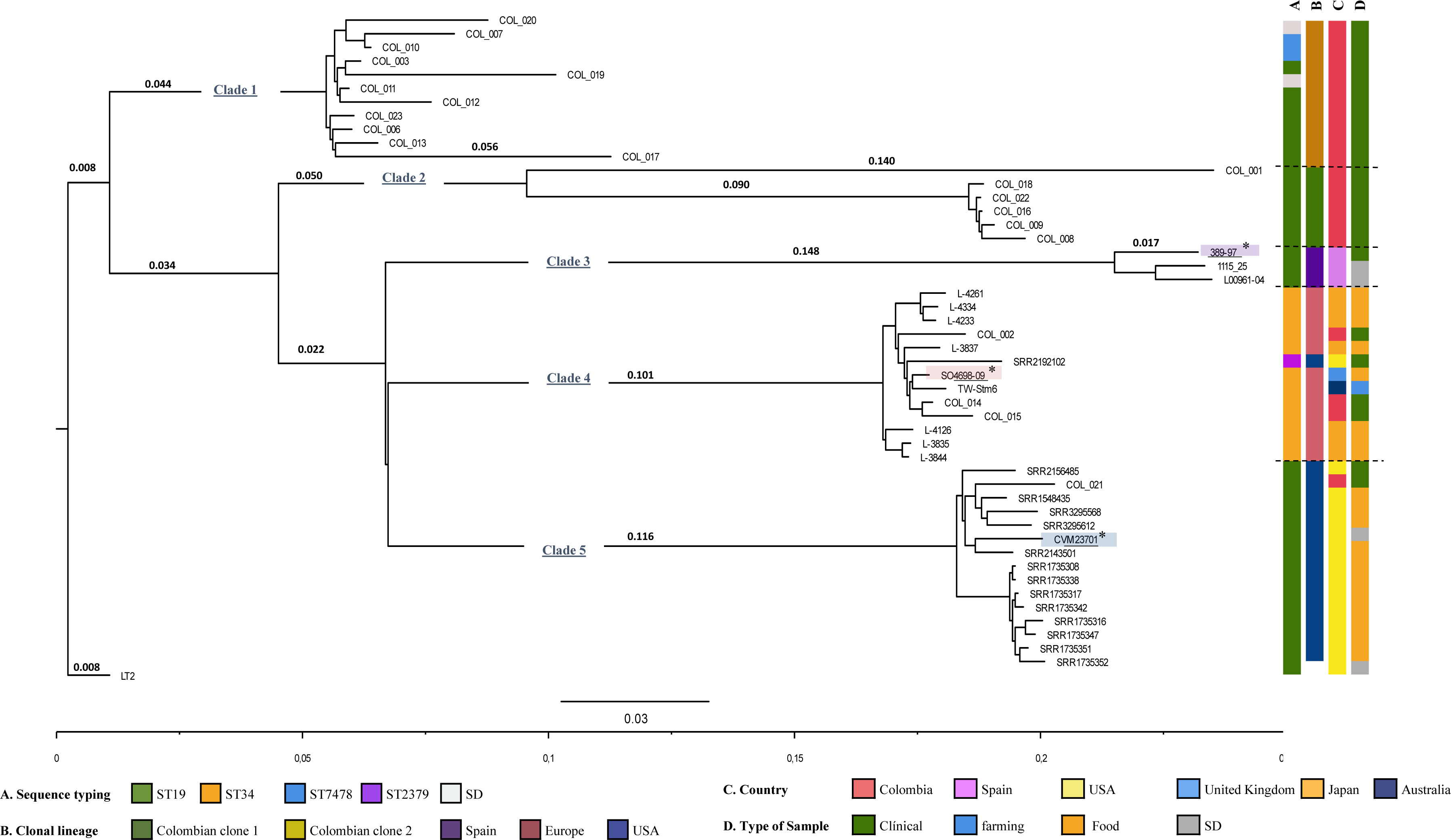
Phylogenetic analysis of Colombian and international STVM isolates. Dendrogram generated by maximum likelihood from SNPs. STVM isolates recovered from 1940 to 2018 from 6 different countries are distributed in 5 clades, a clear differentiation between Colombian isolates clades 1 and 2 from international isolates is observed. Reference strains are highlighted with an * and colored underline in the following order: Purple: Spanish clonal lineage, Salmon: European clonal lineage and Blue: US clonal lineage. The columns to the right indicate a color code found at the bottom of the figure: A) sequencing, B) clonal lineage, C) country of origin and D) sample type.

The phylogeny was constructed from the concatenated alignment of high-quality SNPs for 48 genomes from animal, environmental, and clinical sources in Colombia, the UK, the USA, Australia, Spain, and Japan. Five clades were identified: clades 1 and 2 included 80% (n=17) of the Colombian genomes, while clades 3, 4, and 5 included the international lineages. Clade 3 included isolates classified as the Spanish ST19 clone, clade 4 included the European ST34 lineage, and clade 5 included the USA ST19 lineage. Of the Colombian genomes, only 3 showed a relationship with the European MVST ST34 clone (COL_002, COL_014, COL_015) and 1 with the USA MVST ST19 lineage (COL_021) (Figure 1). These results suggest that the Colombian MVST isolates could be part of two endemic clones and are not related to the clones circulating internationally. The data were analyzed using the microreact platform with the following access link: https://microreact.org/project/9YzKMfHLqSMT1VH4BgALP2-comparativegenomicsmvst

To estimate the introduction or emergence of MVST in Colombia, a Bayesian analysis was performed. Five clusters defined by the most recent common ancestor (MRCA) are evident, preserving the clade grouping of the phylogenetic tree (Figure 2). It is estimated that the MRCA of cluster 1 appeared in 2006, giving rise to a possible Colombian clone 1 that expanded and became established between 2011 and 2012, while cluster 2 shares an MRCA with the international clones approximately since 1964 and stabilized in 2011, giving rise to Colombian clone 2. These results confirm the circulation of 2 Colombian endemic clones from 2011. The estimated dates of appearance for the ST34 European, ST19 Spanish, and ST19 USA lineages in the Bayesian analysis agree with those reported in the literature (32).

**Figure 2.**
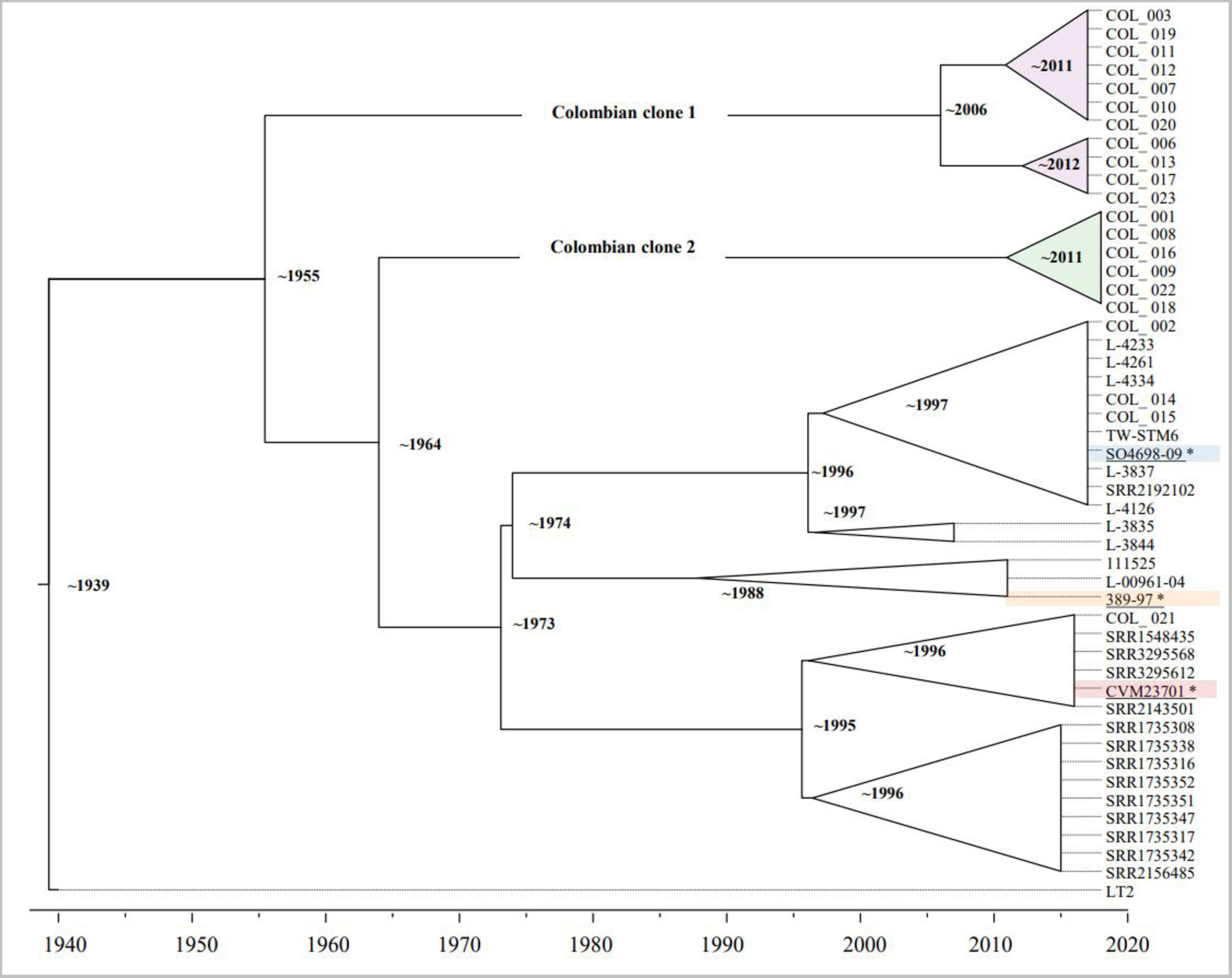
Time phylogeny analysis of Colombian vs. international MVST isolates. Maximum likelihood phylogeny according to Bayesian information criterion (BIC) to understand the phylodynamic of MVST samples and estimate divergence dates. A distribution like the phylogenetic tree is evidenced, confirming the circulation of 2 Colombian lineages highlighted in purple and green, whose clonal expansion is estimated between the years 2011-2012. The estimated dates of appearance for the international lineages are located in the 1990s (consistent with what is reported in the literature). Reference strains are highlighted with an * and colored underline in the following order: European clonal lineage, Spanish clonal lineage, Spanish clonal lineage and USA clonal lineage.

### Colombian lineages are variable in the pattern of chromosomal deletions

One of the conserved characteristics that identify MVST isolates is the deletion of the region between the STM2762 and STM2773 genes, with variations in some isolates. In 12 of 21 Colombian MVST genomes, a complete deletion was observed from gene STM2765 to the last gene of the *fljAB* operon. The remaining 9 isolates partially or completely conserved the *fljAB* operon genes and the *hin* gene: isolate COL_003 conserved *fljB*, COL_006 conserved fljA, COL_007 conserved *fljB* and *hin* genes; COL_008, COL_009, COL_016, COL_018, and COL_022 conserved *hin* and isolate COL_001 conserved the three *fljAB* operon genes although phenotypically it did not express the second flagellar phase (Figure 3). In Colombian clone 1 the STM2762 and STM2763 genes are conserved in all isolates, while they are absent in clone 2 (except COL_009). Regarding the genes upstream of the operon, the presence of *iroB* is conserved in almost all isolates regardless of clones.

**Figure 3.**
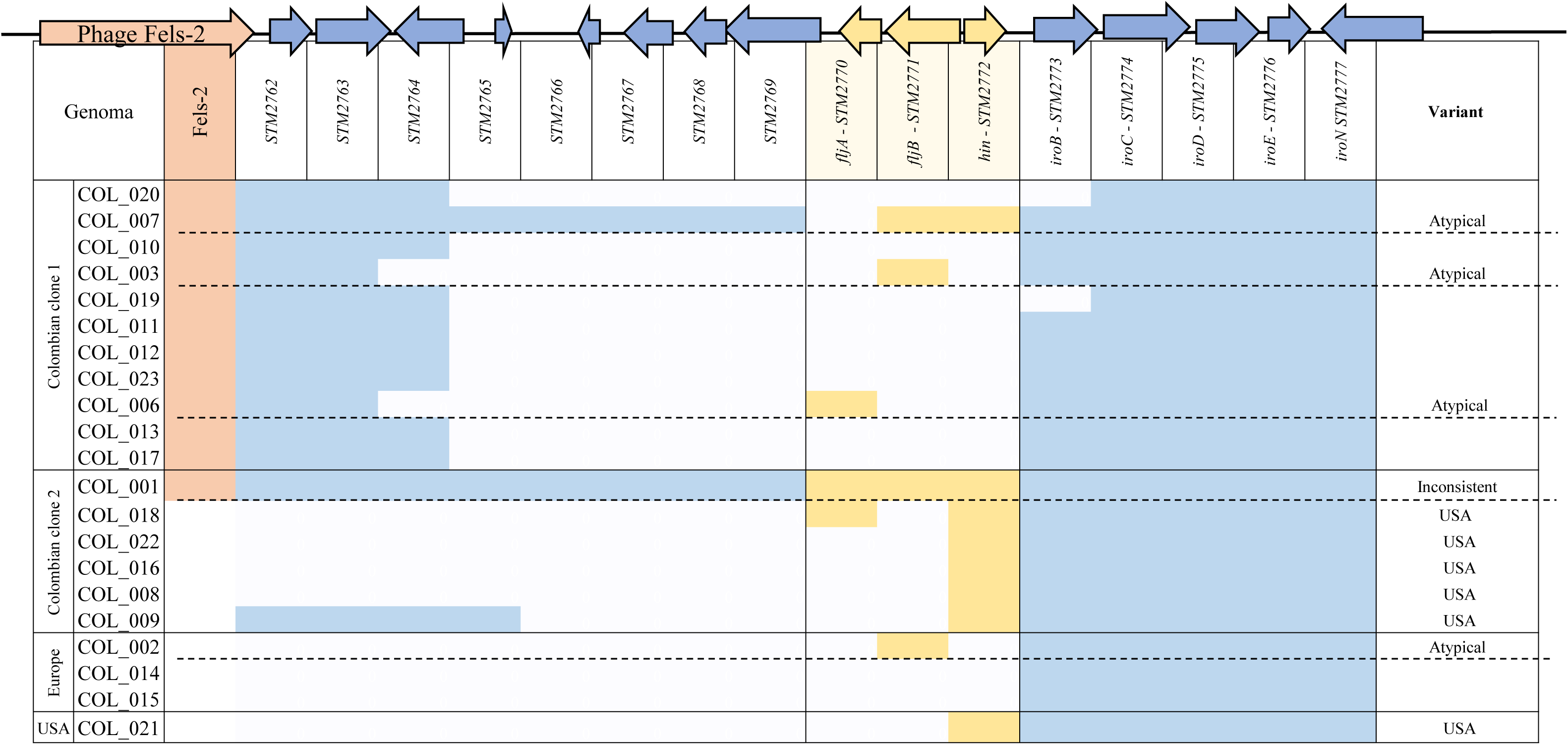
Pattern of deletions in the genomic environment of the *fljAB* operon. Deletions of the *fljAB* operon and its adjacent regions were compared in the 21 Colombian isolates, 17 isolates lack the *fljAB* operon, a pattern of deletions is observed between clonal lineages of genes downstream of the *fljAB* operon and loss of the upstream *iroB* gene in 4 isolates. Colored regions indicate presence and white absence of the gene. The salmon region indicates the presence of the Fels-2 prophage, the yellow region corresponds to the 3 genes of the *fljAB* operon, and the blue colored regions indicate the genomic environment of the *fljAB* operon. On the right side of the figure is the preliminary classification based on the pattern of deletions of the *fljAB* operon, Atypical variant: retains one or two genes of the operon; Inconsistent variant: retains all three genes of the operon, although phenotypically it only exhibits the phase 1 flagellar antigen; USA variant: only retains *hin*.

Upon complete alignment of the MVST sequences against the Typhimurium LT2 reference strain, it was observed that the deletion patterns by a genome were random; however, by clonal lineages, some deletions in the same region of the chromosome are shared (Supplementary Figure 1). The absence of the prophage characteristic *Salmonella* spp. as a determining feature in the evolution of the phase 2 negative variant is discussed below.

We particularly found the deletion of 8 genes (STM3812 to STM3819) in 81% (n=17/21) of the Colombian MVSTs, corresponding to the *ccmABCDEFGH* operon (duplicated operon on the Typhimurium chromosome) responsible for cytochrome C maturation, which is present in all 27 international isolates analyzed.

### The Colombian MVST clones harbor different plasmids, antibiotic-resistance genes, and SGI genomic islands

Thirteen incompatibility groups of virulence or resistance plasmids were identified. Colombian genomes carry from 1 to 11 incompatibility groups. Incompatibility groups related to the two conformations of the Typhimurium virulence plasmid (pSTV) were identified: wild pSTV (IncFIB) and hybrid pSTV (IncFII-IncFIB), 37% (n=18/48) of the genomes conserved the wild pSTV (n=3 genomes Colombian clone 1, n=1 Colombian clone 2), 31% (n= 15/48) presented the hybrid pSTV, 13 Colombian genomes (in both clones) and 3 from the USA. This result is confirmed by the identification of genes exclusive to the Typhimurium virulence plasmid, such as the virulence operon *spvRABCD,* the fimbrial genes *pefABCD*, and the *rck* gene, in the genomes with the presence of the IncFII and IncFIB groups. The 3 genomes (14.3%) grouped with the European ST34 clone lack this pSTV as well the genes associated to this plasmid (Table supp 2).

Additionally, a greater number of incompatibility groups were observed in Colombian clone 1, and Colombian clone 2 exhibits fewer incompatibility groups as can be seen in Table 1, where COL_009 isolate MDR carries 8 of the described groups (Figure 4).

**Figure 4.**
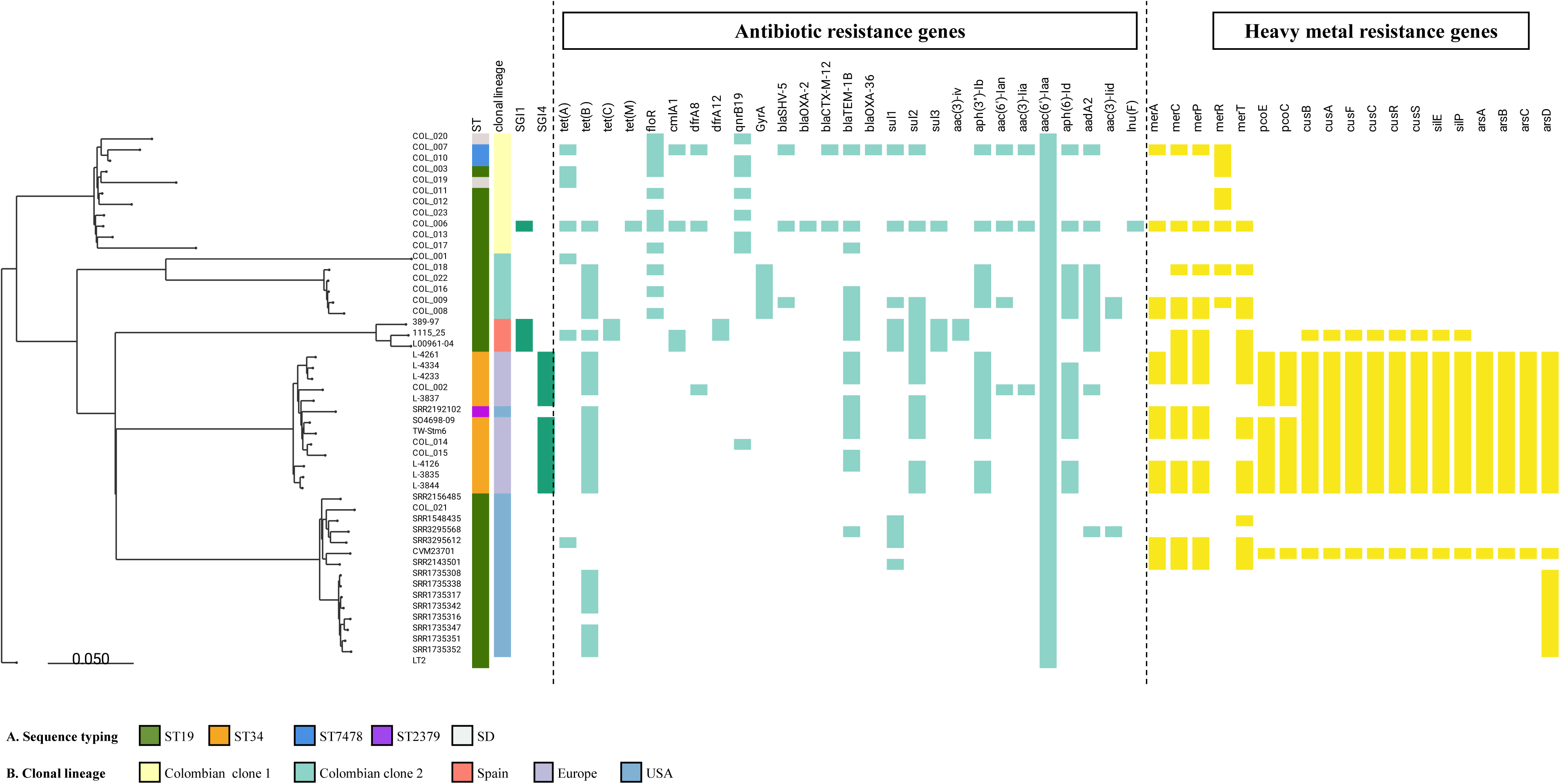
Acquisition of genetic elements of antimicrobial and heavy metal resistance. Resistance profiles in Colombian and international MVST are shown next to the dendogram. The columns to the right indicate a color code found at the bottom of the figure: A) sequencing, B) clonal lineage. The presence of two genomic islands SGI-1 and SGI-2 is indicated in green, resistance to 5 antibiotic families in light blue and resistance to heavy metals in yellow.

**Table 1.** Incompatibility groups were observed in Colombian clones. The plasmid incompatibility groups for the Colombian clones are grouped according to the number of genomes (n) that present them and their respective percentage.

| Incompatibility groups | Colombian clone 1<br>(n=11) |  | Colombian clone 2<br>(n=6) |  |
| --- | --- | --- | --- | --- |
|  | n | % | n | % |
| IncI1-I(Alpha) | 9 | 82 | 3 | 50 |
| ColRNAI | 5 | 45 | 1 | 17 |
| Col(VCM04) | 1 | 9 | 3 | 50 |
| Col(pHAD28) | 4 | 36 | 1 | 17 |
| ColpVC | 3 | 27 | 1 | 17 |
| IncQ1 | 0 | - | 2 | 33 |
| IncC | 2 | 18 | 0 | - |
| IncFIA | 2 | 18 | 0 | - |
| IncHI1B(R27) | 2 | 18 | 0 | - |
| IncA/C2 | 2 | 18 | 0 | - |

The IncI1-I(Alpha) plasmid was identified in isolate Col_007 with an approximate size of 92,000 base pairs, 95.9% similarity percentage, and 82% coverage with that reported by Oliva et al 2020 (accession number MT507877) in a clinical monophasic isolate ST1030 recovered in southern Italy in 2008 (33). In this plasmid the replication and transfer modules are mainly conserved, the aligned contigs show the mercury resistance operon *merRTPCADES* and although no contig is observed splicing with the region composed of the Tn21 carrying the resistance genes such as *aadA1-cmlA1-aadA2*, in isolates COL_ 007 and COL_006 we found the *cmlA1* and *aadA2* genes, but *aadA1* is absent.

This incompatibility group was identified in 9 genomes of clone 1 and 3 genomes of clone 2. The coverage percentages were between 82% and 21% in 10 genomes, which indicates that this genetic element is characteristic of the Colombian clones, except in COL_023 in which only one contig was identified with this Inc group but no additional contigs were found that matched the reference plasmid ST1030. The *mer* operon was one of the major segment differences absent in all 10 IncI1-I(Alpha) plasmids (Table Supp3).

A total of 24 resistance genes were identified in the Colombian genomes, belonging to 8 antibiotic families as shown in (Figure 5). Resistance to chloramphenicol mediated by *floR* is found in both Colombian clones (n=11), as well as resistance genes to penicillins, cephalosporins and aztreonam: *bla*SHV-5 (n= 3), *bla*TEM-1B (n= 7) and sulfonamides: *sul1* (n= 3), *sul2* (n=4).

**Figure 5.**
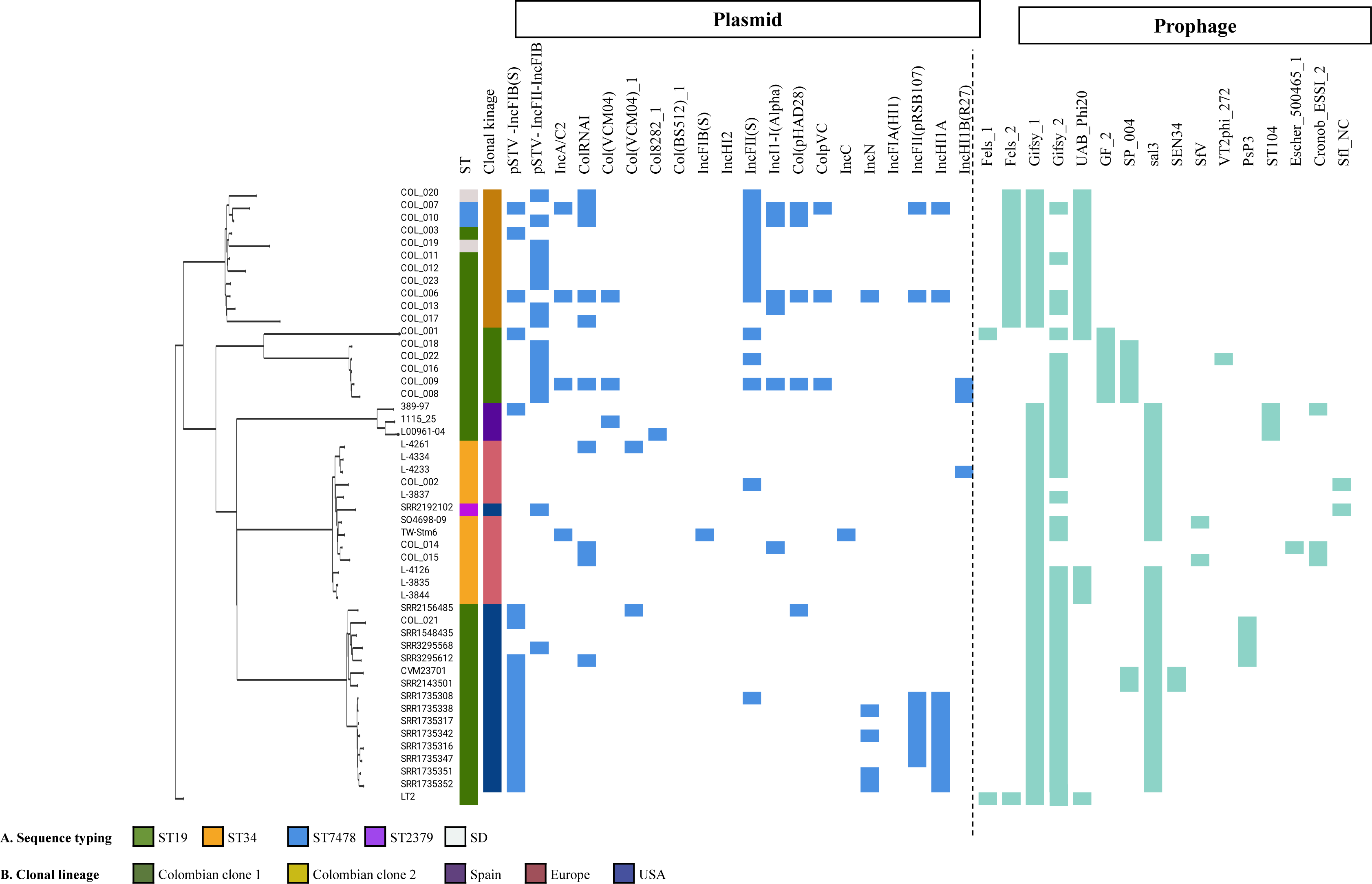
Acquisition of plasmids and prophages. The acquisition of plasmids and the integration of prophages in Colombian and international MVSTs are shown next to the Dendrogram. The columns to the right indicate a color code found at the bottom of the figure: A) sequencing, B) clonal lineage. Plasmid incompatibility groups are shown in dark blue and prophage regions in light blue.

In the Colombian clone 1, there are 21 resistance genes, of which 9 seem specific for this clone, as shown in Table 2, including a resistance gene to lincosamide *(inuF)*. This last gene reported in Gram-positive bacteria is present in isolate COL_006 that is multidrug resistant (MDR) and carries Genomic Island I (SGI-1) responsible for resistance to 7 antibiotics: ampicillin, chloramphenicol, florfenicol, streptomycin, spectinomycin, sulfonamides and tetracycline (34).

**Table 2.**
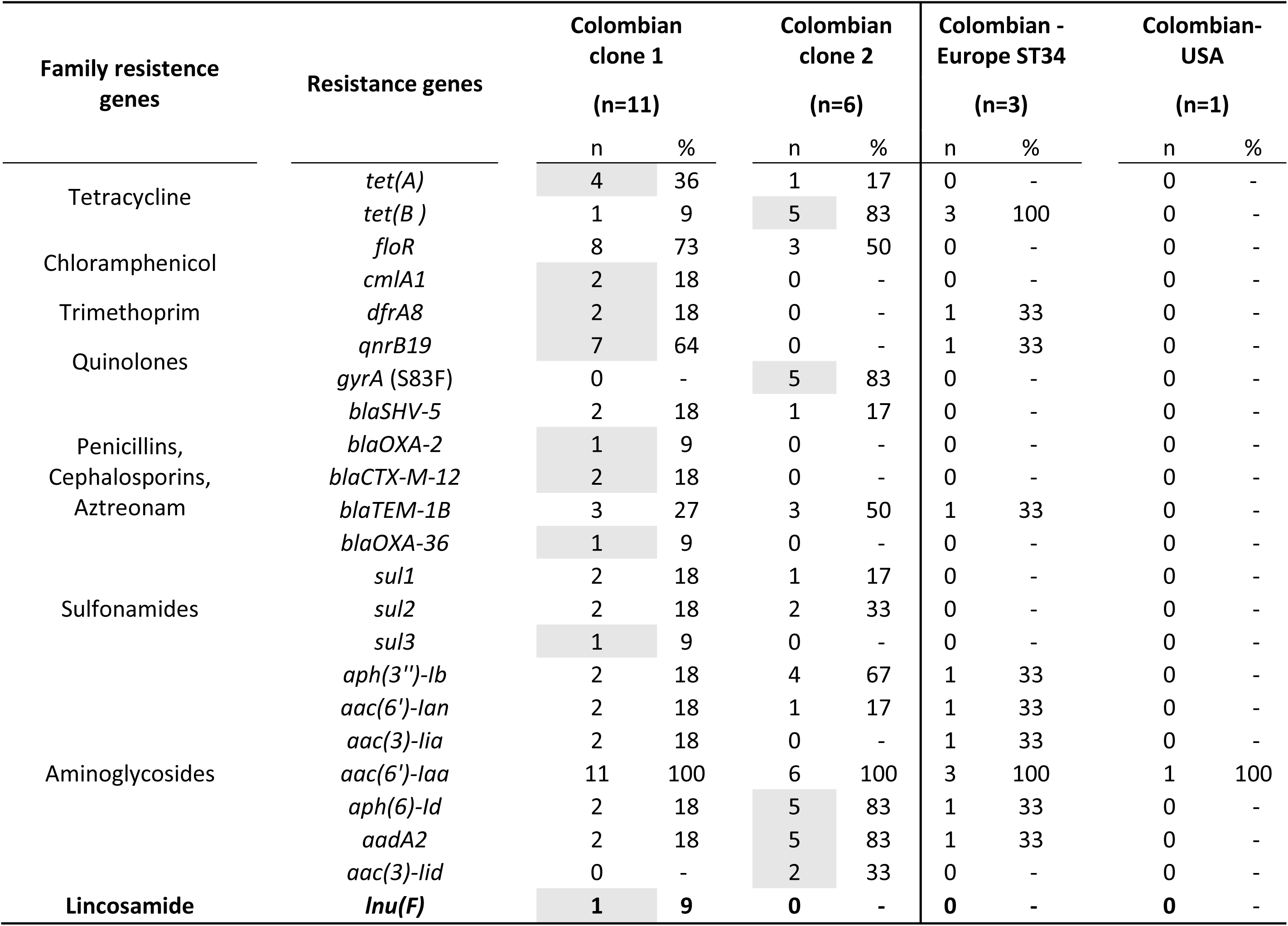
Resistence genes in Colombian clones The table shows a total of 24 resistance genes for 8 families of antibiotics, the Colombian genomes were distributed according to their phylogenetic relationship, are grouped according to the number of genomes (n) that present them and their respective percentage. The genes associated with Colombian clones 1 and 2 are highlighted. The lincosamide resistance gene is reported in a darker font.

The Colombian clone 2 shows patterns of resistance to antimicrobials and heavy metals similar to those reported in international clones, its main characteristic is the presence of *tet(B)* that codes for resistance to tetracyclines as well as resistance to quinolones due to the S83F mutation in the gene *gyrA* (Table 2).

Antimicrobial susceptibility confirmed in the laboratory showed a correlation between the genes identified and the antimicrobial resistance expressed in each isolate. We observed that the predominant resistance in the two Colombian clones is tetracycline (100%), nalidixic acid (66.6%) and chloramphenicol (57.1%) (https://microreact.org/project/9YzKMfHLqSMT1VH4BgALP2-comparativegenomicsmvst).

Although in the genomes COL_006 and COL_007 of Colombian clone 1 and COL_008, COL_009 and COL_018 of Colombian clone 2 the *merRTPCADES* operon conferring mercury resistance was identified, it appears to be inserted in different genomic contexts. In the genomes of clone 1 and COL_009 of clone 2, the *mer* operon is inserted in the plasmid of the IncI1-I group described in the previous section. In the other two genomes, it was not possible to define whether this element is inserted in these or another plasmid or the chromosome. However, the results show that this element is mostly present in Colombian clone 2 (3 genomes out of 5). It was also observed that the first gene of the operon which is *merR* is found in the COL_003, COL_010, COL_011, and COL_ 012 genomes, but the remaining genes of the operon are absent (Supplementary figure 2).

The genomes COL_002, COL_014, and COL_015, which belong to the European clone are the only ones containing genomic island 4 (SGI-4), with genes coding for tolerance to heavy metals such as Copper (*pcoGEABCDRSE*), Silver (*silESRCFBAGP*) and Arsenic (*arsRDABC*) (Figure 4).

The lack of Typhimurium-specific prophages is characteristic of Colombian clone 2. In general, all MVST clusters analyzed in this study show the absence of the Typhimurium-specific prophages Fels-1 and Fels-2. The previously reported international clonal specificities were confirmed in this analysis. In total, 16 prophage regions were identified in the 48 genomes analyzed. In the international clones grouped in clusters 3, 4, and 5, 12 of the 16 described prophages were identified. In common, all carry Gifsy_1, 92% (n=25/27) carry the Sal3 prophage and 85% (n=23) carry Gifsy_2. The Colombian genomes grouped in the Europe clone ST34 lack these two prophages. In the Spanish clone ST19 (cluster 3) additionally, the ST104 prophage and a genome with the SfI_NC prophage were identified. In the European clone ST34 and the USA clone ST19, UAB_Phi20, Cronob_ESSI_2, SfV, Escher_500465_1, SfI_NC, SP004, PsP3 and SEN34 were additionally identified (Figure 4 and Table 3). In the Colombian clone, the prophage repertoire is different among them and concerning the international clones, carrying 8 of the 16 identified regions. In Colombian clone 1, 100% (n=11) of the genomes conserve the Fels_2, Gifsy_1, and UAB_Phi20 prophages, and four genomes carry the Gifsy_2 prophage, while Colombian clone 2 exhibits the most variable prophage profile, with the exclusive presence of GF_2 in 100% (n=6), followed by Gifsy_2 and the acquisition of SP004 in 83% (n=5) of the genomes, with both found in 4 genomes simultaneously. As for particular prophages the COL_001 genome is the only one that conserves Fels_1 and COL_016 presents VT2phi_272 (Figure 4). All the Colombian genomes were negative for the mTmV phage, a characteristic element of the European clone ST34 and ST19.

**Table 3.** Prophage were observed in Colombian clones The table shows the distribution of the prophages in the Colombian genomes, these were distributed according to their phylogenetic relationship, they are grouped according to the number of genomes (n) and their respective percentage. Prophages associated with each clonal lineage are highlighted.

| Prophage | Colombian clone 1<br>(n=11) |  | Colombian clone 2<br>(n=6) |  | Colombian -Europe<br>ST34<br>(n=3) |  | Colombian- USA<br>ST19<br>(n=1) |  |
| --- | --- | --- | --- | --- | --- | --- | --- | --- |
|  | n | % | n | % | n | % | n | % |
| Fels_1 | 0 | - | 1 | 17 | 0 | - | 0 | - |
| Fels_2 | 11 | 100 | 0 | - | 0 | - | 0 | - |
| Gifsy_1 | 11 | 100 | 0 | - | 3 | 100 | 1 | 100 |
| Gifsy_2 | 4 | 36 | 5 | 83 | 0 | - | 1 | 100 |
| UAB_Phi20 | 11 | 100 | 1 | 17 | 0 | - | 0 | - |
| GF_2 | 0 | - | 6 | 100 | 0 | - | 0 | - |
| SP_004 | 0 | - | 5 | 83 | 0 | - | 0 | - |
| sal3 | 0 | - | 0 | - | 1 | 33 | 1 | 100 |
| SEN34 | 0 | - | 0 | - | 0 | - | 0 | - |
| SfV | 0 | - | 0 | - | 1 | 33 | 0 | - |
| VT2phi_272 | 0 | - | 1 | 17 | 0 | - | 0 | - |
| PsP3 | 0 | - | 0 | - | 0 | - | 1 | 100 |
| Escher_500465_1 | 0 | - | 0 | - | 1 | 33 | 0 | - |
| Cronob_ESSI_2 | 0 | - | 0 | - | 2 | 67 | 0 | - |
| Sfi_NC | 0 | - | 0 | - | 1 | 33 | 0 | - |

### *Salmonella* virulence factors and *Salmonella* pathogenicity islands (SPIs) are conserved elements in MVST genomes

Supplementary table 2, shows the search for virulence factors with two web resources (VFDB and BV-BRC). A total of 146 effector proteins related to virulence and cell survival were found, of which 88 are conserved in the 48 genomes analyzed. In the two Colombian lineages, the presence of the characteristic genes of the pSTV virulence plasmid was identified.

On the contrary, the genomes grouped in the Spanish clone ST19, European clone ST34 and some of the clone ST19 USA where we observe the absence of the fimbrial operon *pefABCD* in 35.4% (16/48) and of the *spvRABCD* operon in 39.5% (19/48), which is in agreement with the reports where it is mentioned that these clones have lost the pSTV. As for chromosomal virulence genes, the *cheABD* operon involved in chemotaxis is absent in most clusters, the *cheA* gene is absent in 96% of the genomes, *cheB* in 94%, and *cheD* in 92%. The most frequently missing virulence factors are *rfbI* (98%), *spaM* (96%), *gtrA* (95%), *manB* (90%), *entF* (86%), and *shdA* (41%). All 48 genomes analyzed have similar virulence factor content and maintain chromosomal content integrity.

SPIFinder identified 12 *Salmonella* pathogenicity islands in the 48 genomes analyzed: SPI-1, SPI-2, SPI-3, SPI-4, SPI-5, SPI-9, SPI-12, SPI-13, SPI-14, Centisome 63 (C63PI), Centisome 54 (CS54) and one putative island. SPI-1, SPI-2, SPI-5, SPI-13, and SPI-14 were detected in 100% of the genomes. SPI-4 is conserved in 77% of the genomes, SPI-9 in 90.5%, C63PI in 47.6% and CS54 in 38.1%. SPI-4 and C63PI are predominant in clone 2, while in clone 1 there are present in 50% and 27% of the genomes respectively, CS54 is present in 3 genomes of each clone. SPI-12 was only found in international isolates related to the European clonal lineage ST34. As for the virulence genes associated with the prophages acquired by the European MVST clones, contrary to expectations, the Colombian genomes do not carry the *sopE* gene of the mTmV phage, nor the *grvA* gene of the Gifsy2 prophage.

## Discussion

The nature of the monophasic variant has allowed its successful dissemination throughout the world, overcoming the Typhimurium serovar in the last decades (7), adapting to different ecological niches, especially the pork food chain and other meat products for human consumption in Europe, USA and China (7, 35, 36), suggesting a direct transmission of the bacterium to humans from the consumption of contaminated products.

The high incidence of MVST (monophasic variant of *Salmonella* Typhimurium) in human salmonellosis cases poses a significant public health risk. This was evident in a recent outbreak caused by the consumption of chocolate products contaminated with this serovar, which affected 10 countries with 150 reported cases, predominantly among children (37).To effectively monitor the spread of clones and evolutionary events, as well as global and local transmission, the implementation of tools such as whole genome sequencing (WGS) is crucial. In Colombia, the Instituto Nacional de Salud (INS) surveillance program has observed a rise in clinical cases of MVST, surpassing the prevalence of the previously dominant serovar Typhimurium, which had been prominent in the country for the past decade (12, 13). Given the global emergence of this serovar, it is important to understand its genetic characteristics and determine whether its successful dissemination is attributed to the introduction and circulation of an international clone or the emergence of an endemic clone. Such knowledge is essential for effective control and prevention measures.

Previous genomic studies conducted on Typhimurium isolates in Colombia between 1997 and 2006 confirmed the presence of two European clones, namely ST34 and ST39, circulating in the country. Phylogenetic analysis also revealed that certain monophasic ST19 isolates were closely related to a Typhimurium isolate recovered from fecal material in the USA, as well as to Typhimurium isolates associated with the LT2 reference strain (15).

In this study, we expanded upon these findings by evaluating 21 Colombian genomes (11 that were previously sequenced). These genomes were collected between 2015 and 2018 and were compared with 27 genomes from five different countries, including reference strains from MVST international clones. The analysis revealed that 80% of the Colombian genomes clustered into two distinct groups, which were separate from the international clones. These groups were designated as Colombian clone 1 and Colombian clone 2. Additionally, we identified the co-circulation of the international European ST34 and USA ST19 clones in five isolates, which had been previously reported in the study by Li et al. in 2019. This finding contrasts with our previous studies, in which we employed PCR characterization by amplifying the *fljAB* operon region (14). By utilizing whole genome sequencing, which is a more accurate and reliable methodology, and by conducting a phylogenetic comparison with reference clone sequences, we can affirm that the Colombian clinical MVST isolates belong to an endemic lineage that became established in the country in 2011. The analysis of chromosome deletion patterns, one of the initial characteristics identified in the evolution of the MVST variant (38), did not reveal a conserved pattern among the 21 Colombian genomes analyzed. However, we did observe significant differences in the accessory genome between the two clones, which also differed from the international clones. These findings suggest a local adaptation of the clinical isolates of MVST in Colombia, highlighting the influence of the local environment on the genetic makeup of these isolates.

Genomic analyses have provided evidence of the global circulation of three major clones: the Spanish clone ST19, the European clone ST34, and the USA clone ST19 (7), These clones have also served as the basis for the emergence of recent endemic lineages, as demonstrated by the study conducted by Ingle et al (2021) in Australia (39). Regarding the Colombian lineage, the variation in its accessory genome plays a crucial role in distinguishing between two clones within a small number of isolates. The results revealed the presence of two distinct groups of accessory genomes, each corresponding to one of the identified clones. A detailed explanation of these findings is provided below.

Clone 1 is characterized by the presence of 9 Inc groups, including pSTV, multiple resistance plasmids, predominantly IncI1-I(Alpha), and several ColRNA1 type plasmids. Within this clone, there are genes conferring resistance to tetracycline (*tetA*), fluoroquinolones (*qnrb19*), various aminoglycosides (*aadA*), as well as *sul1*, *sul2*, and *sul3* genes associated with sulfonamide resistance. Furthermore, multiple beta-lactam resistance genes (*bla*) are also present. It has been documented in the literature that these genes are commonly carried within the Inc groups identified in this study, often found within class 1 integrons and transposons. Additionally, previous investigations conducted by our research group have demonstrated that the *tetA* gene in Colombian Typhimurium isolates is mainly located within the ΔTn1721 element, which is carried by plasmids belonging to the IncF, IncA/C, and IncX1 families (unpublished results, https://www.ins.gov.co/Direcciones/Investigacion/Proyectos/Caracterizaci%C3%B3n%20de%20los%20determinantes%20gen%C3%A9ticos%20asociados%20a%20la%20multirresistencia%20en%20aislamientos%20cl%C3%ADnicos%20de%20Salmonella%20Typ.pdf).

Clone 2 exhibits a lower number of Inc groups compared to Clone 1, as well as a reduced presence of resistance genes. In contrast to Clone 1, fluoroquinolone resistance in Clone 2 is conferred by a mutation in the chromosomal gene *gyrA*. Similarly, tetracycline resistance in this clone is attributed to the presence of the *tetB* gene, which is inserted within the ΔTn10 element, found to be located in both plasmids and chromosomes in Typhimurium Colombian isolates (unpublished results, https://www.ins.gov.co/BibliotecaDigital/Caracterizacion-determinantes-geneticos-asociados-multirresistencia-aislamientos-cl%C3%ADnicos-Salmonella-Typhimurium.pdf).

An intriguing finding is the presence of the Fels-2 prophage in Clone 1, which is absent in the international clones. It is speculated that this prophage has been replaced by the mTmV prophage carrying the virulence gene *sopE*, which is also lacking in the Colombian isolates (32). Unlike the conserved Gifsy prophages found in monophasic clones, the Fels-1 and Fels- 2 prophages are typically absent in monophasic strains worldwide (20, 38, 40). The Fels-2 prophage, typically associated with Typhimurium LT2 (41), is rarely encountered in other Typhimurium lineages. Therefore, the identification of the conserved Fels-2 prophage in Colombian Clone 1 suggests a closer genetic relationship with Typhimurium LT2 or a potential descent from a biphasic LT2-like ancestor (42). In the case of Clone 2, it exhibits the most diverse phage profile compared to the LT2 reference strain. It carries phages such as SP_004, whose presence has been described in Typhimurium isolates from sources like dairy cattle and dairy products in the USA (6). Additionally, it possesses the GF-2 phage, specific to *Edwardsiella tarda* (43). The content genes of GF-2 possess lysogeny-related genes that have not been found in the other reported *Edwardsiella* phages and by comparative genomics of *Edwardsiella myophages* it is suggested that the C-terminal domains of the tail fiber proteins of this phage have relevance to its host specificity (44). It has been reported that virulence gene transfer can be responsive to and dependent on variations in bacterial environmental conditions and help explain the emergence of new *Salmonella* outbreaks through the acquisition of new prophages.

This data set suggests that clone 1 has an accessory genome more enriched in mobile genetic elements such as plasmids, transposons, and integrons than clone 2.

These results correlate with those reported in the analyses of population-based studies in MVST (45), which recently demonstrated that MVST sequences from different countries are mainly grouped in a cluster 1 that also mostly conserves a particular accessory genome. Ingle et al. (2021) reports the circulation of three Australian lineages of MVST, each associated with different resistance genes and different deletions in the *fljAB* operon (39). The European ST34 lineage is primarily characterized by the acquisition of the SGI4 island (46). These observations suggest that the evolution and adaptation of MVST isolates are largely influenced by the acquisition of accessory genomes rather than variations in their core genome. However, it should be noted that the core genomes of MVST variants with different geographical origins do exhibit changes. In the Colombian clones, the chromosomal deletions are not uniform, indicating genomic plasticity in the microevolutionary process. It remains unclear whether these deletions have been replaced by transposons. These findings indicate that the spread of the monophasic variant in Colombia is not solely attributed to a Salmonellosis outbreak but rather reflects the genomic plasticity of MVST within the country. Additionally, our observations reveal that the Colombian MVST lineage lacks the second copy of the *ccmABCDEFGH* operon, which is responsible for cytochrome c maturation. While cytochrome c maturation systems are known to be independent, it is noteworthy that system I can mature cytochromes C at heme concentrations five times lower than system II and can function as a "heme reservoir," suggesting an adaptation to survive in different environments (47, 48). The biological implications of the absence of this operon in Colombian MVST isolates remain unknown.

One notable characteristic observed in the Colombian clones is their resistance to a smaller number of antibiotic families compared to the international clones. Consequently, they carry fewer resistance genes, except for MDR genomes that also harbor the *mer* operon. Interestingly, these isolates possess the IncI1 plasmid, which shares 82% similarity to a previously reported MDR clinical isolate recovered in Italy (49).In contrast to the Colombian clones, Chinese isolates predominantly carry IncHI2-IncHI2A plasmids, as observed in previous studies. The Australian lineage is characterized by the predominance of IncA/C, IncI, and IncHI2 plasmids, while the Japanese isolates are associated with the pYT1 plasmid, a hybrid virus-resistance plasmid (39, 50).

These findings highlight the variations in resistance plasmids among MVST isolates from different regions, where the IncI1 plasmid is predominantly carried by the Colombian clones. Although the IncI1 plasmid is not the predominant element circulating in Colombia, it appears to be associated with isolates of serovar Typhimurium and MVST (15) . The presence of the *qnrB* gene, conferring resistance to fluoroquinolones and typically encoded in small plasmids of the ColRNAI type (15), is particularly noteworthy. Alongside the IncI1 plasmid, these elements, predominantly associated with clone 1, contribute to antibiotic resistance against agents that have been classified by the World Health Organization (WHO) as high-priority targets in combating antimicrobial resistance. The presence of these elements may provide a selective advantage to clone 1 over clone 2 in MVST isolates, as the presence of mercury resistance also serves as a selection pressure in natural and production environments (51).

The Colombian lineages 1 and 2 in the country have emerged through separate introduction and dissemination events. Among them, Colombian lineage 1 exhibits greater similarity to the reference strain and has adapted to the environment by acquiring diverse plasmids and antibiotic resistance genes. These genetic elements are likely associated with human hosts, where the selective pressure in clinical environments facilitates the transfer of these genes between bacteria of different species. Consequently, MVST serves as a reservoir for genes like *inuF*, although its acquisition does not provide an adaptive advantage for *Salmonella*. On the other hand, Colombian lineage 2 bears resemblance to the USA clone, incorporating prophages such as SP_004, which are associated with pig fattening farms in the USA. This suggests that the introduction of this clonal lineage to Colombia may have occurred through the importation of meat products, sharing MRCA with international lineages. Subsequently, it underwent a differentiation process and adapted to the Colombian population.

These findings demonstrate the remarkable plasticity of MVST and underscore the importance of active genomic surveillance to monitor the circulation of new clonal lineages in Colombia. Such monitoring is crucial as these lineages have the potential to significantly impact public health.

### Ethics approval and consent to participate

This research has the approval of the Technical Research Committee (CTIN) and the Ethics Research Committee (CEIN) of the Instituto Nacional de Salud (INS), Colombia with the codes CTIN-05-2015 and CTIN-05-2017. Clinical and patient ID from whom the MVST clinical isolates were recovered, were anonymized.

## Acknowledgements

We acknowledge all public health laboratories from the national network of laboratories. To the Genome sequencing performed at the Earlham Institute as part of the 10KSG consortium which is supported by the Global Challenges Research Fund data and resources grant (BBS/OS/GC/000009D). Next-generation sequencing and library construction were delivered via the BBSRC National Capability in Genomics and Single Cell (BB/CCG1720/1) at Earlham Institute, by members of the Genomics Pipelines Group. To Diego Prada for advice in bioinformatics tools.

## Financials

No external funding was received for this study.

---

**Supplementary Figure 1.**
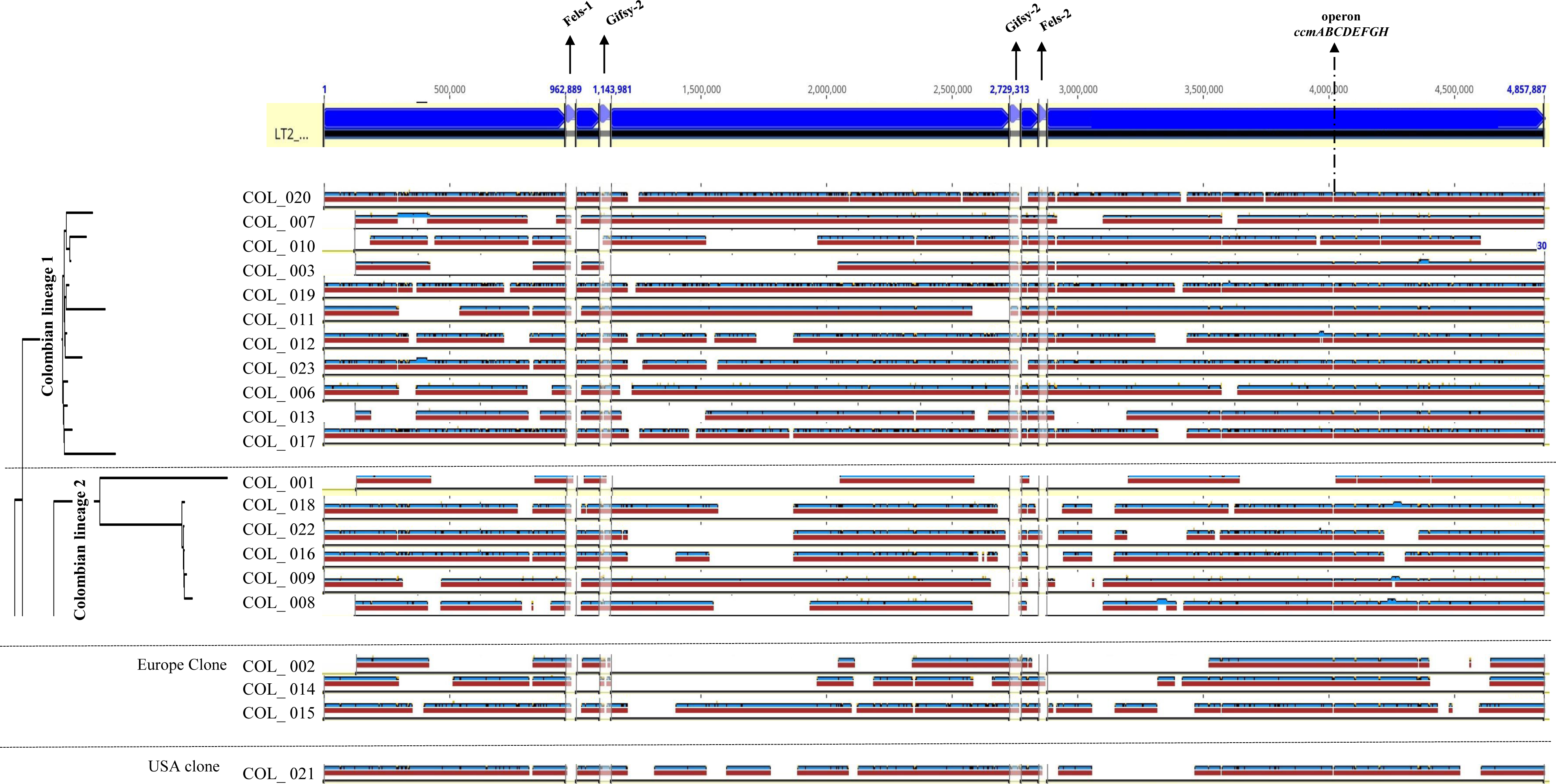
Chromosomal comparison of the LT2 reference strain and Colombian MVST. The figure shows the genomic plasticity of the Colombian MVST isolates according to the clonal groupings obtained by phylogeny. In the upper part of the figure is the LT2 chromosome with a size of 4.8Mb, the smaller arrows demarcated by black lines represent the regions of the LT2 prophages: Fels-1, Gifsy-2, Fels-2, Gifsy-2. On the right side, the deletion site of the *ccmABCDEFGH* operon (duplicated operon on the Typhimurium chromosome) is indicated by a dotted line.

**Supplementary Figure 2.**
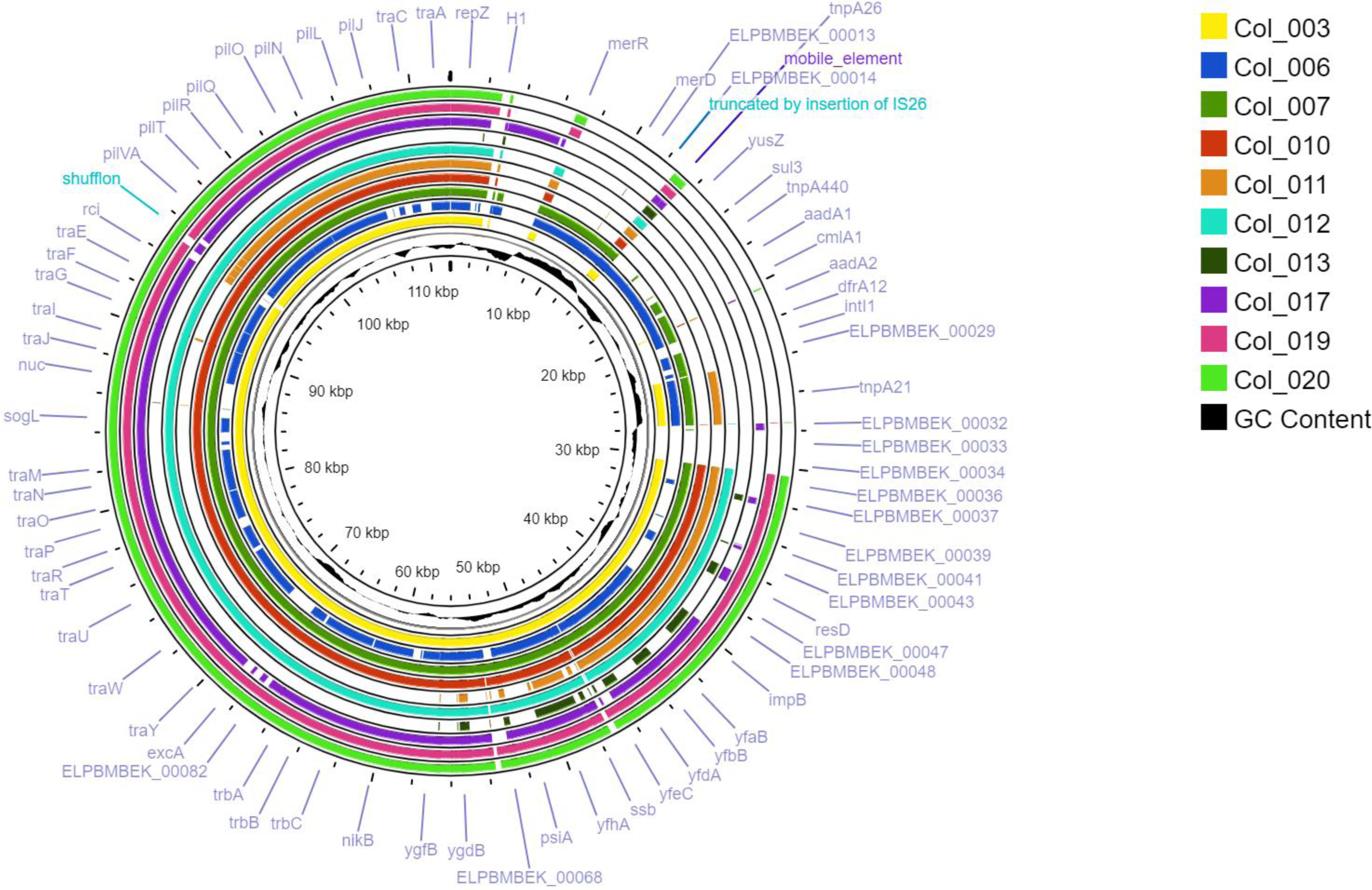
Alignment of IncI1 MT507877 plasmid regions found in 10 genomes of Colombian clone 1. On the right side of the figure, we can observe the color coding for the respective genomes. The genomes present a great similarity, however, they lack the variable region where an int1 would be found.

**Table supplementary 3.** Identification of plasmids with IncI1 (alpha) Incompatibility group Result of the search and mapping of plasmids with IncI1 (alpha) incompatibility group, where the reference plasmid, the percentage of identity and the percentage of coverage among the Colombian genomes are related.

|  | Genome | Incompatibility<br>group IncI1 (alfa) | Plasmid_ref | Plasmid name | pb ref_plasmid | % identity | Base of<br>pairs (pb) | % coverage |
| --- | --- | --- | --- | --- | --- | --- | --- | --- |
| Colombian<br>clone 1 | Col_020 | IncI1 (alfa) | MT507877 | pST1030-1A | 112.136 | 95,8 | 87927 | 78.4110366 |
|  | Col_007 |  |  |  |  | 95,8 | 92650 | 82.62288649 |
|  | Col_010 |  |  |  |  | 95,2 | 89206 | 79.55161589 |
|  | Col_003 |  |  |  |  | 95,4 | 89551 | 79.85927802 |
|  | Col_019 |  |  |  |  | Identical Sites: 23456<br>(99.9%) | 23480 | 20.93885996 |
|  | Col_011 |  |  |  |  | Identical Sites: 42021<br>(90.7%) | 46261 | 41.25436969 |
|  | Col_012 |  |  |  |  | 95,2 | 89152 | 79.50346008 |
|  | Col_023 |  |  |  |  | Identical Sites:3700<br>(92.8%) | 3982 | 3.551045159 |
|  | Col_006 |  |  |  |  | 94,3 | 69320 | 61.81779268 |
| Colombian<br>clone 1 | Col_001 | IncI1 (alfa) | MT507877 | pST1030-1A | 112.136 | Identical Sites: 43569<br>(97.3%) | 44712 | 39.87301134 |
|  | Col_022 |  |  |  |  | Identical Sites: 3700<br>(92.8%) | 3982 | 3.551045159 |
|  | Col_009 |  |  |  |  | Identical Sites: 50353<br>(91.8%) | 54693 | 48.77381037 |

